# Leaf hydraulics is a core component of plant immunity

**DOI:** 10.64898/2026.07.27.740543

**Authors:** Soline Marty, Caroline Bellenot, Damien Vasselon, Magali Goussot, Alice Boulanger, Corinne Audran, Pauline Savourat, Anne-Sophie Sarthou, Cécile Pouzet, Jean-Marc Routaboul, Patrick Laufs, Laurent D. Noël

## Abstract

Hydathodes at leaf margins mediate guttation of xylem-derived fluids and serve as primary entry sites for adapted vascular bacterial pathogens such as *Xanthomonas campestris*. Infection of Arabidopsis mutants with fewer hydathodes resulted in spontaneous mesophyll water-soaking and revealed unexpectedly-large pathogen populations explained by direct infection of the mesophyll niche through stomata and subsequent proliferation. Physical blockage of hydathodes also induced mesophyll water-soaking and enhanced bacterial growth in both Arabidopsis and cauliflower leaves. These findings reveal a dual role for hydathodes as primary sites of infection while being essential to restrict pathogen proliferation in nonvascular tissues. More broadly, our study identifies leaf hydraulics and guttation as central components of water immunity in vascular plants.

## Main Text

Water is essential for all living organisms, including plant pathogens, which must acquire water from their hosts during infection. To this end, pathogens have evolved diverse strategies to increase humidity within their niche in the leaf apoplast, thereby facilitating bacterial motility and diffusion of nutrients required for growth (*1*). These processes often culminate in the appearance of dark-green, water-soaked leaf lesions caused by fluid accumulation in apoplastic spaces before visible tissue collapse. Under normal growth conditions, apoplastic water accumulation (also called leaf flooding) is rarely observed and occurs mostly upon infection.

Several physiological mechanisms can promote water-soaking, including increased ambient humidity, altered stomatal behavior that modulates vapor exchange, and changes in vascular water transport (*2*). Phytopathogenic bacteria can suppress stomatal immunity, the closure of stomatal triggered by pathogen-associated molecular pattern (PAMP) perception, and thus promote water accumulation in leaves (*3*). For instance, coronatine, a jasmonic acid mimic produced by *Pseudomonas syringae* pv. *tomato* (*Pst*), suppresses stomatal closure and thereby promotes infection. However, mutant plants defective in stomatal closure display enhanced resistance to *Pst* infection because increased desiccation of the apoplast restricts pathogen proliferation, a process called water immunity (*4*). These findings underscore the central role of water regulation in plant immunity and further indicate that water immunity is potent as it can compensate for defects in stomatal immunity. Beyond stomatal regulation, water immunity may also involve reduced local vascular activity through decreased xylem conductivity, as well as reduced abundance or activity of aquaporins (*5, 6*).

Consistent with the importance of water immunity, several type III effector (T3E) proteins injected by phytopathogenic bacteria promote water-soaking and contribute to disease development. For instance, WtsE T3E protein enhances both water accumulation and the release of metabolites in the apoplast of maize infected by *Pantoea stewartii* subsp. *stewartii* (*7*). AvrHah1, produced by X*anthomonas gardneri*, induces the release of oligosaccharides, thereby increasing extracellular sugar availability and driving water influx into infected tissues (*8*). In *Pst*, AvrE and HopM1 T3E proteins were identified as key redundant determinants of apoplastic hydration, disease progression, and immune suppression (*9, 10*). These effectors interfere with multiple host pathways, including the vesicle-trafficking regulator AtMIN7, the PP2A phosphatase involved in activation of anti-PAMP immune receptor complexes, and the abscisic acid (ABA) transporter ABCG40, which promotes stomatal closure following infection (*11-14*). Altogether, such T3E functions evidence the key role of access to water for pathogens during their interaction with plants.

Hydathodes, located at leaf margins in vascular plants, are other key regulators of leaf water status. These organs are directly connected to xylem vessels and mediate guttation, the release of xylem-derived fluid when evapotranspiration is limited (*15*). Hydathode pores resemble enlarged stomata that respond to light and ABA but do not fully close (*16*). Beneath these pores lies a loosely organized parenchyma named epithem connected to a hypertrophied xylem vasculature. Hydathodes function as hydraulic pressure-release valves that protect the leaf from water-soaking. Disruption of the bundle sheath in *Arabidopsis thaliana* mutants abolishes guttation and causes mesophyll water-soaking under conditions that normally promote guttation (*17*). Likewise, mechanically sealing hydathodes or reducing their abundance such as in the *cuc2* and *sty* mutants result in enhanced tissue water-soaking (*18, 19*). Hydathodes provide a nutrient- and water-rich environment (*20*) which represent preferred entry sites for several vascular pathogens, including *Xanthomonas campestris* pv. *campestris* (*Xcc*), the causal agent of black rot disease in Brassicaceae (*21*). After entering through hydathodes, *Xcc* colonizes xylem vessels and spreads systemically. Notably, *Xcc* cannot initiate infection through stomata, even in mutants with constitutively open stomata (*16*). In contrast, the closely related nonvascular pathogen *Xanthomonas campestris* pv. *raphani* (*Xcr*) infects via stomata and causes leaf spot disease. These distinct infection strategies may reflect major differences in the water status of the tissues encountered by bacteria during colonization. For instance, increased water availability alone can promote the proliferation of bacteria nonpathogenic on plants, highlighting access to water as a major determinant of pathogen success (*22-24*).

In this study, we establish an essential role for hydathodes in determining *Xcc* ecological niche during the early stages of infection and demonstrate that guttation and leaf hydraulics are core components of water immunity in vascular plants.

### Reduced number of hydathodes promotes spontaneous water-soaking associated with ectopic proliferation of *Xcc*

To investigate the genetic basis of hydathode immunity in *Arabidopsis thaliana* Col-0, we screened a random subset of 366 lines from the Homozygous EMS Mutant (HEM) collection (*25*) for altered bacterial proliferation of the virulent *Xcc* strain 8004Δ*avrAC* in the rosette (*26, 27*). The HEM collection comprises sequenced mutant lines carrying an average of 576 mutations per genome, ca. 85% of which are homozygous (*28*). To favour hydathode infection, plants were spray-inoculated and maintained under high humidity (fig. 1a, fig. S1a). Bacterial populations were quantified from whole rosettes at eight days post-inoculation (dpi). This screen identified mutant lines displaying either enhanced or reduced bacterial proliferation relative to the Col-0 parental line and the susceptible accession Oy-0 (fig. S1b, data S1) (*29*). Because hydathodes constitute the primary entry sites for *Xcc*, we counted the number of hydathodes on the third leaf of each line (fig. 1b, data S1). Unexpectedly, the number of hydathodes was not positively correlated with an increased bacterial load across the mutant population. Even more surprisingly, it showed a negative and weak, yet non-significant, correlation with bacterial load (ρ = –0.092, p-value = 0.08), suggesting that reduced hydathode number might, if at all, favor proliferation rather than limit it (fig. 1b). To investigate this paradox, we analyzed mutants and transgenic lines with altered numbers of hydathodes. These included mutations affecting the transcription factors CUP-SHAPED COTYLEDON 2 (CUC2), which regulates leaf serration and hydathode development (*30-32*), and DORNRÖSCHEN (DRN) and DORNRÖSCHEN-LIKE (DRNL), which function in organ primordia and auxin signaling pathways associated with hydathode formation (*33-35*). Strong *cuc2-1* mutants and *drn-1 drnl-1* double mutants exhibited markedly reduced hydathode numbers, whereas *CUC2* overexpression (*CUC2g-m4*) approximately doubled the number of hydathodes relative to wild type (fig. 1c, fig. S2). Despite having fewer hydathodes, *cuc2-1* and *drn-1 drnl-1* plants accumulated ca. 10-fold higher *Xcc* populations than wild type, whereas *CUC2g-m4* and the weak *cuc2-3* allele showed no increase in bacterial load (fig. 1c). These results support the negative trend observed in the HEM population and unexpectedly indicate that hydathodes are not limiting for infection. Instead, fewer hydathodes may open alternative infection routes into leaf tissues. To localize bacterial proliferation sites, plants were inoculated with *Xcc* 8004Δ*avrAC::GUS-GFP*. At six dpi, GUS staining remained largely restricted to hydathodes in wild-type, *cuc2-3*, and *CUC2g-m4* plants, whereas *cuc2-1* and *drn-1 drnl-1* mutants displayed extensive colonization of the mesophyll (fig. 1d). Mesophyll colonization correlated with the appearance of spontaneous water-soaking during the dark period, consistent with impaired guttation caused by reduced hydathode number (fig. 1e). Similarly, spontaneous water-soaking mutants of *CURLY LEAF clf-28* and *clf-29* (*36*) developed severe disease symptoms and supported extensive proliferation of *Xcc* 8004Δ*avrAC::GUS-GFP* in the mesophyll (fig. S3). Taken together, these results suggest that mesophyll water-soaking could be necessary and sufficient to promote ectopic proliferation of *Xcc*.

**Fig. 1.**
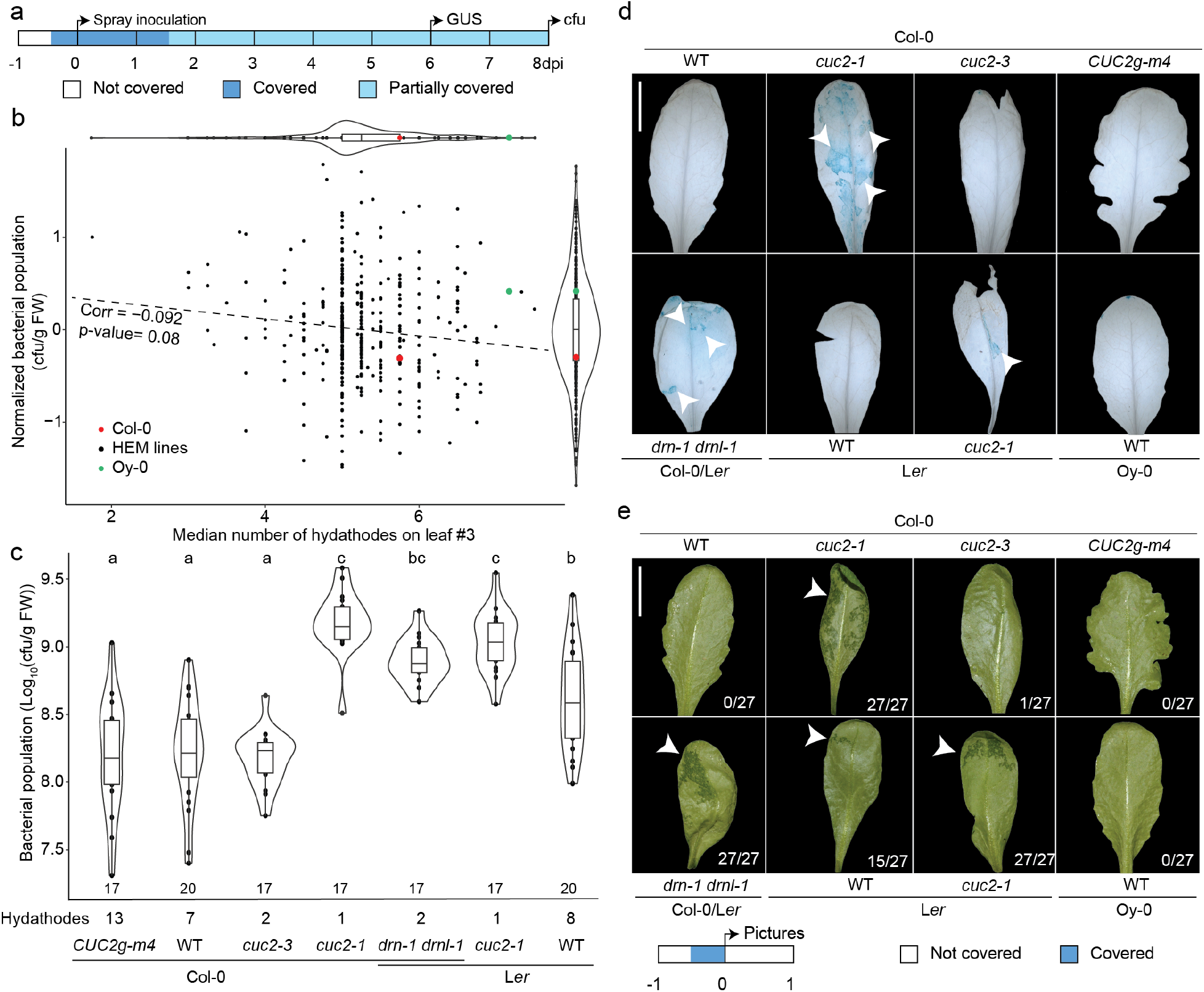
Hydathodes limit leaf spontaneous water-soaking and associated pathogen proliferation in the mesophyll of Arabidopsis leaves. **(a)** A timeline depicting the inoculation procedure and sampling is shown. cfu = colony forming unit, dpi = days post inoculation. **(b)** Scatter plot representation of the normalized *Xcc* 8004*ΔavrAC* populations in the whole rosette at eight dpi and the number of hydathodes on the third leaf in 366 HEM mutant lines and wild-type Col-0 parental line and the more susceptible accession Oy-0. Each point represents the values for the median bacterial population and the median number of hydathodes for one HEM line (Data S1). Three plants were phenotyped in three biologically-independent experiments. The Spearman correlation (p = 0.08) is represented by a dashed line. Data distribution along each axis is shown using violin plots. **(c)** Boxplot representation of *Xcc* 8004*ΔavrAC* populations at eight dpi in *A. thaliana* mutants and transgenic lines and corresponding wild types showing variable numbers of hydathodes on the fifth leaves. Letters indicate statistically distinct groups according to a linear model with adjusted comparison (p<0.05). The number of leaves sampled to determine the bacterial population is indicated. **(d)** GUS activity produced by *Xcc* 8004*ΔavrAC::GUS-GFP* at six dpi in *A. thaliana* mutants, transgenic lines and corresponding wild types. Arrowheads indicate *Xcc* proliferation in the mesophyll tissue. **(e)** Spontaneous water-soaking of mesophyll (arrowheads) is observed in *cuc2-1* and *drn-1 drnl-1* mutants that have a reduced number of hydathodes after one night in high humidity conditions. Numbers indicate the frequency of leaves showing water-soaking. Scale bar = 1 cm.

### Guttation prevents mesophyll water-soaking and restricts stomatal colonization by *Xcc*

To directly test whether guttation restricts mesophyll colonization by *Xcc* and to bypass any pleiotropic developmental defects that may be present in the mutants used above, we phenocopied hydathode-deficient mutants by sealing hydathodes at leaf margins with silicone grease. Blocking guttation induced pronounced mesophyll water-soaking by the end of the dark period under high humidity growth conditions (fig. 2a,b). Following dip inoculation with the virulent strain *Xcc* 8004Δ*avrAC::GUS-GFP*, bacterial proliferation was detected within substomatal chambers exclusively in sealed leaves and as early as one dpi (fig. 2c, fig. S4). By 2-3 dpi, *Xcc* had reached high population densities throughout the adjacent mesophyll apoplast, whereas virtually no proliferation was observed in nonsealed leaves. These observations indicate that blocking guttation enables *Xcc* to enter through stomata and initiate infection from substomatal cavities. This again demonstrates that hydathode function is essential to prevent mesophyll colonization by *Xcc*.

**Fig. 2.**
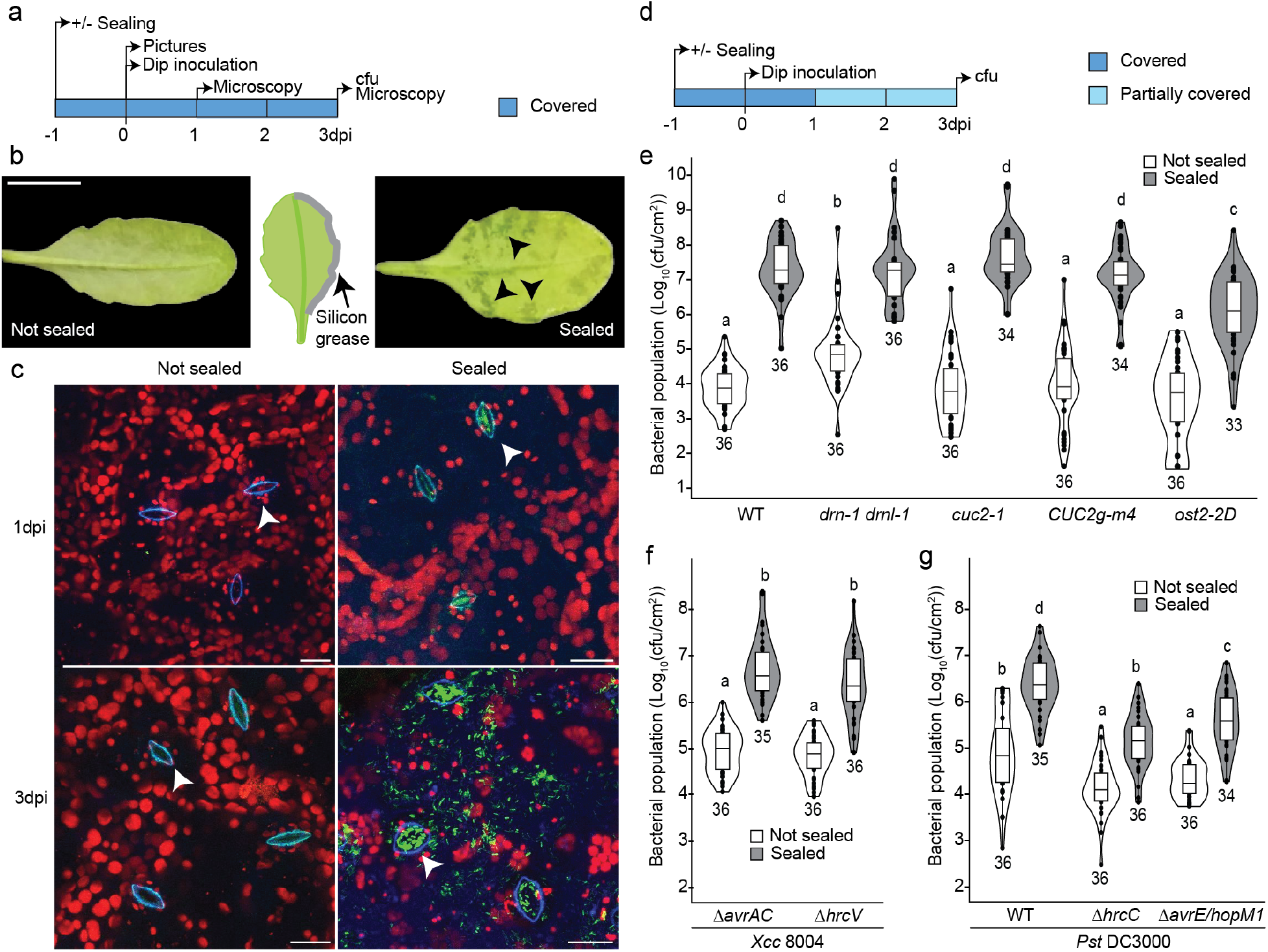
Mechanical suppression of guttation in Arabidopsis leaves allows for bacterial proliferation in stomatal chambers of the mesophyll. **(a**,**d)** Timelines depicting the inoculation and sampling procedures are shown for b,c and e-g, respectively. **(b)** Sealing leaf margins of Col-0 with silicone grease causes water-soaking (arrowheads) 24h after application. Scale bar = 1 cm. **(c)** Observation of abaxial faces of Col-0 leaves by confocal microscopy at 1 or 3 days post dip inoculation with *Xcc* 8004*ΔavrAC::GUS-GFP* with or without sealing the margins. GFP fluorescence (green) can be observed in substomatal chambers in sealed conditions only. Arrowheads indicate stomata. Scale bar = 20 µm. **(e)** Boxplot representation of *Xcc* 8004*ΔavrAC* populations in *A. thaliana* mutants and transgenic lines and corresponding wild types showing variable numbers of hydathodes (*cuc2, drn-1 drnl-1, CUC2g-m4)* or constitutively-opened stomata (*ost2-2D*) with or without sealing leaf margins at three dpi. **(f)** Boxplot representation of bacterial populations of the virulent *Xcc* 8004*ΔavrAC strain* or the avirulent *Xcc* 8004*ΔhrcV* in Col-0, with or without sealing leaf margins at three dpi by dipping. **(g)** Boxplot representation of bacterial population of *Pst* DC3000, *Pst* DC3000Δ*hrcC* or *Pst* DC3000Δ*avrE/hopM1* in Col-0, with or without sealing at three dpi. Letters indicate statistically distinct groups according to a linear (e) or linear mixed-effects model (f,g) with adjusted comparison (p<0.05). The number of leaves sampled is indicated.

To quantify this effect, we measured bacterial proliferation in the mesophyll of wild-type (Col-0), mutant (*cuc2-1, drn-1 drnl-1, ost2-2D*), and transgenic (*CUC2g-m4*) plants with or without leaf-margin sealing (fig. 2d,e). The *ost2-2D* mutant carries constitutively open stomata that fail to close fully in response to environmental stimuli(*37*). In nonsealed conditions, only *drn-1 drnl-1* plants displayed elevated bacterial loads relative to Col-0 and L*er*, consistent with earlier observations (fig. 2e, fig. 1c, fig. S5). By contrast, *cuc2-1* plants were not yet significantly different from wild type at three dpi in contrast to what is later observed at eight dpi (fig. 1c). At three dpi, leaf-margin sealing consistently caused a strong increase in bacterial populations across all genotypes, yet to a lower extent in *ost2-2D* mutant (fig. 2e). This indicates that elevated stomatal conductance partially compensates for the loss of guttation and that *Xcc* entry into leaves at stomata is not sufficient to allow its proliferation. In contrast to our observation in the nonsealed condition, the bacterial load in *drn-1 drnl-1* was identical to the wild type. These results support the conclusion that apoplastic water availability is a major determinant of *Xcc* proliferation within the mesophyll. Under the same high-humidity conditions, the nonpathogenic type III secretion mutant *Xcc* 8004Δ*hrc*V proliferated to levels comparable to those of the virulent strain 8004Δ*avrAC* (fig. 2f), indicating that elevated humidity can bypass the normally essential immune-suppressive functions of type III effectors.

In order to bypass stomatal immunity, direct infection of the mesophyll by *Xcc* was used and yielded similar results as observed using dip inoculation (fig. S6a,b). Only a weak difference between sealed and nonsealed conditions was observed in *ost2-2D* plants. However, under nonsealed conditions, *drn-1 drnl-1* plants supported higher bacterial populations than wild-type plants, suggesting that infiltration-derived water is sufficient to promote *Xcc* proliferation in this mutant background. In *drn-1 drnl-1* plants, sealing did not promote bacterial proliferation suggesting that apoplastic water levels were already sufficient for *Xcc* in nonsealed condition. Disease symptoms closely mirrored bacterial growth patterns. Together, these results establish guttation as critical to restrict *Xcc* proliferation to hydathodes only and reveal the importance of apoplastic water in the mesophyll to control *Xcc* proliferation.

### Guttation also restricts the proliferation of a mesophyll-adapted bacterial pathogen

In contrast to the vascular pathogen *Xcc*, which is normally restricted to hydathodes, *Pst* naturally infects leaves through stomata. This mesophyll-adapted pathogen promotes apoplastic water accumulation through toxins and type III effectors, including AvrE and HopM1, which suppress water immunity and enhance disease development. We therefore asked whether hydathode function also contributes to resistance against *Pst* in *Arabidopsis*. Wild-type *Pst* DC3000 and two mutants unable to cause water-soaking DC3000Δ*hrcC* (type III secretion-deficient) and DC3000Δ*avrE/hopM1* (type III effector mutant) were inoculated onto sealed and nonsealed leaves by dip inoculation (fig. 2g). Sealing leaf margins increased bacterial proliferation for all strains. Under nonsealed conditions, Δ*hrcC* and Δ*avrE/hopM1* mutants accumulated to levels approximately 10-fold lower than those of wild-type DC3000. Remarkably, sealing restored growth of both mutants, with DC3000Δ*hrcC* reaching bacterial populations comparable to those of the nonsealed wild-type strain and DC3000Δ*avrE/hopM1* exceeding them. However, DC3000Δ*avrE/hopM1* did not reach wild-type levels in sealed leaves. Similar trends were observed following direct mesophyll inoculation by infiltration (fig. S6c). Leaf-margin sealing enhanced bacterial proliferation for all strains, whereas the Δ*avrE/hopM1* mutant displayed intermediate growth between wild type and Δ*hrcC* mutant. Thus, loss of AvrE and HopM1 was partially compensated by blocking guttation. Together, these results demonstrate that guttation is important for water immunity established against mesophyll-adapted pathogens.

### Hydathodes and ambient humidity are key components of water immunity in cauliflower leaves

To determine whether hydathode-dependent water immunity extends beyond *Arabidopsis* model plants, we investigated the role of hydathodes in resistance to *Xcc* in the cauliflower crop from which strain 8004 was isolated (*38*). Plants were maintained either covered or noncovered throughout the infection period to manipulate ambient humidity, and a subset of leaves was additionally sealed at the margins to inhibit guttation and increase apoplastic water accumulation (fig. 3a). Intense water-soaking was observed in the sealed leaves (fig. S7a). Leaves were dip-inoculated with the virulent *Xcc* 8004::*GUS-GFP* strain or the avirulent type III secretion mutant 8004Δ*hrcV*. By three dpi, severe disease symptoms developed specifically in sealed leaves inoculated with the wild-type strain, whereas leaves inoculated with the Δ*hrcV* mutant or mock-treated leaves remained essentially asymptomatic (fig. 3b). Nonsealed leaves showed little or no visible disease symptoms. GUS staining revealed that bacterial proliferation remained largely restricted to hydathodes in nonsealed leaves (fig. S7b). In contrast, sealed leaves displayed extensive mesophyll colonization by the wild-type strain, particularly under covered conditions (fig. 3c, fig. S7b). Notably, detectable mesophyll proliferation of the Δ*hrcV* mutant was also observed in sealed and covered leaves, indicating that elevated apoplastic water availability can partially compensate for the absence of type III secretion. Quantification of bacterial populations and co-localization of water-soaked areas with infection confirmed that ambient humidity and apoplastic water accumulation additively promoted bacterial proliferation for both strains (fig. 3d, fig. S7c). Together, these results demonstrate that hydathode function and environmental humidity cooperate to establish water immunity and restrict *Xcc* proliferation within the cauliflower mesophyll.

**Fig. 3.**
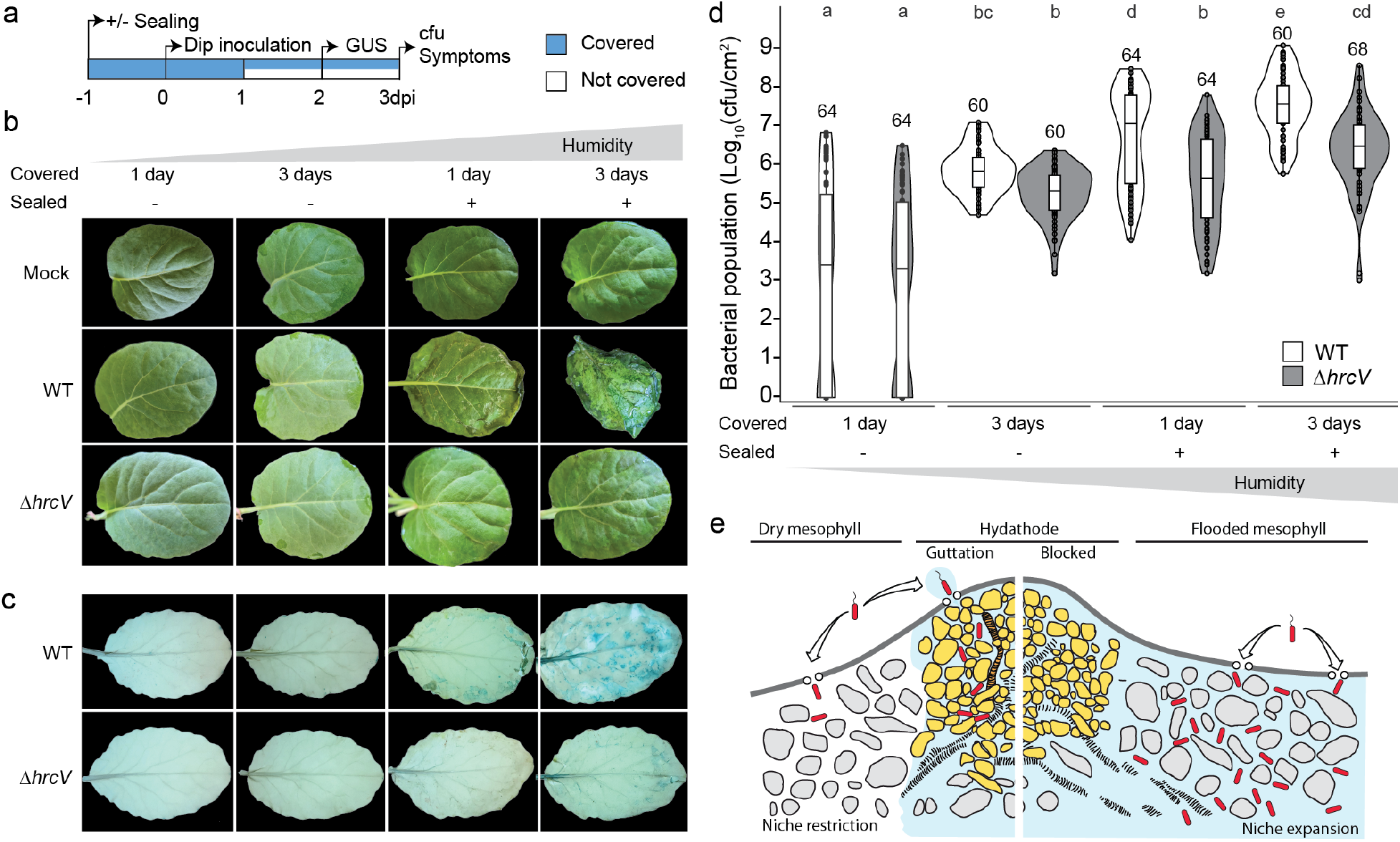
Promotion of virulent and avirulent *Xcc* population on cauliflower by increasing water-soaking and ambient humidity. **(a)** A timeline depicting the inoculation procedure and sampling is shown. **(b)** Symptoms caused by virulent *Xcc* 8004 (WT) and avirulent *Xcc* 8004*ΔhrcV* (*ΔhrcV*) strains on the second true leaf of cauliflower leaf depending on humidity and sealing of leaf margins at three days post dip inoculation. **(c)** GUS activity produced by *Xcc* 8004*::GUS-GFP* (WT) or *Xcc* 8004*ΔhrcV::GUS-GFP* (*ΔhrcV*) reporter on cauliflower leaves at two dpi. **(d)** Boxplot representation of population of virulent *Xcc* 8004 (WT) and avirulent *Xcc* 8004*ΔhrcV* (*ΔhrcV*) in cauliflower depending on environmental humidity and sealing at three dpi. Letters indicate statistically distinct groups according to a linear mixed-effects model with adjusted comparison (p<0.05). The number of leaves sampled is indicated. **(e)** Graphical summary illustrating the role of guttation at hydathodes to prevent apoplastic water accumulation (in blue) and promote water-soaking of the mesophyll (grey cells) and proliferation of bacterial pathogens (in red). The epithem is shown in yellow. Representative images are shown.

## Discussion

Guttation and water-soaking are highly dynamic physiological processes which lack proper physiological and genetic characterization. Because hydathodes and guttation are conserved across vascular plants, impaired guttation likely imposes a substantial fitness cost. This cost has traditionally been attributed to reduced gas exchange caused by excessive water accumulation within leaves. Here, we reveal that hydathodes also play defining roles in the establisment of water immunity in the leaves of vascular plants (fig. 3d).

Water-soaking is a hallmark of many bacterial diseases and has generally been considered a consequence of infection (*2, 39*). For instance, *Pst* is able to actively promote water-soaking during infection using phytohormone mimics or type III effector proteins (*10*). Our results demonstrate that sufficient mesophyll hydration is a prerequisite for bacterial proliferation. Both vascular and mesophyll-adapted pathogens proliferated extensively when apoplastic water accumulation was experimentally increased, even in the absence of a functional type III secretion system. Under sufficiently humid conditions, avirulent strains of *Xcc* and *Pst* reached bacterial populations comparable to those of virulent strains in *Arabidopsis* or cauliflower. These observations reveal that restriction of apoplastic water availability constitutes a major layer of plant immunity. This effect was particularly striking for *Xcc*, a vascular pathogen that is normally excluded from the mesophyll at early stages of infection, including in mutants with constitutively open stomata (*16*). Our findings suggest that the mesophyll environment is normally too dry to support efficient *Xcc* proliferation and that hydathode-mediated guttation contributes to maintaining this restrictive state. Consistent with this interpretation, increased transpiration in the *ost2-2D* mutant partially suppressed the effects of experimentally induced water-soaking. Together, these results identify apoplastic water availability as a central determinant of disease outcome and establish water immunity as a core component of plant defenses. Whether similar mechanisms operate in nonvascular plants remains an open question.

The transition between vascular and nonvascular lifestyles represents a major evolutionary shift in plant-associated bacteria. Current models propose that ancestral *Xanthomonas* species were vascular pathogens and that repeated transitions toward mesophyll colonization occurred through independent events of gene gain and loss (*40*). Our findings suggest that adaptation to the mesophyll may critically depend on the ability to manipulate or tolerate low-humidity apoplastic environments. In this context, comparison between vascular *Xcc* strains and closely related nonvascular pathogens such as *Xcr* may help identify the genetic innovations required for adaptation to the for mesophyll niche. These may include mechanisms promoting apoplastic hydration, tolerance to water limitation, or enhanced survival under fluctuating osmotic conditions.

More broadly, our study identifies leaf hydraulics and guttation as central determinants of plant immunity and pathogen niche specialization. By controlling apoplastic water availability, hydathodes define whether leaf tissues remain resistant or permissive to bacterial colonization. Water availability therefore emerges not simply as a consequence of infection, but as a primary ecological and immune barriers shaping plant–pathogen interactions.

## Acknowledgments

We wish to thank Harrold van den Burg (University of Amsterdam, Netherlands) and Peter Moffett and Charles Roussin-Léveillée (University of Sherbrooke, Canada) for sharing seeds of the Oy-0 accession and *clf* mutants, respectively.

## Funding

ANR NEPHRON ANR-18-CE20-0020-01 (CB, DV, PS, AB, ASS, MG, PL, LDN) LABEX TULIP ANR-10-LABX-41 (SM, CB, AB, CA, CP, JMR, LDN) EUR TULIP-GS ANR-18-EURE-0019 (SM, CB, AB, CA, CP, JMR, LDN). EUR Saclay Plant Sciences ANR-17-EUR-0007 (DV, MG, PS, ASS, PL) IDEX Saclay Plant Sciences ANR-11-IDEX-0003-02 (DV, MG, PS, ASS, PL)

## Author contributions

Conceptualization: PL, JMR, LDN

Investigation: SM, CB, DV, PS, MG, ASS, JMR, AB, CA, CP

Data analysis and drafting of the manuscript: SM, LDN

Project administration: PL, LDN

Supervision: PL, LDN Funding acquisition: PL, LDN

Writing – original draft: SM, LDN Writing – review & editing: all authors

## Competing interests

Authors declare that they have no competing interests.

## Data, code, and materials availability

All data are available in the main text or the supplementary materials.

## Notes

### Competing Interest Statement

The authors have declared no competing interest.

