## Supplemental figures and Table for "Leaf hydraulics is a core component of plant immunity"

**The PDF file includes:**

**Fig. S1.** Screening of 366 HEM Arabidopsis mutants for their susceptibility to virulent *Xcc* upon spray inoculation and hydathode-mediated infection.

**Fig. S2.** Observation and counting of hydathodes in leaves of Arabidopsis mutants and transgenics and their corresponding wild-types.

**Fig. S3.** *Xcc* can colonize the mesophyll of *clf* mutants

**Fig. S4.** Observation of abaxial sides of Col-0 leaves by confocal microscopy at 1, 2 or 3 days post dip inoculation with *Xcc* 8004*ΔavrAC::GUS-GFP* with or without leaf margins sealing

**Fig. S5.** Susceptibility of *drn-1 drnl-1* mutant upon dip inoculation with virulent *Xcc 8004ΔavrAC* strain

**Fig. S6.** water-soaking in sealed condition promotes bacterial population after infiltration in Arabidopsis Col-0 leaves

**Fig. S7.** Inhibition of guttation induces water-soaking and mesophyll infection in cauliflower

**Table S1.** List of strains used in this study

**Other Supplementary Materials for this manuscript include the following:**

**Data S1.** List of HEM lines and observed phenotypes

### Materials and Methods

#### Plant material and growth conditions

*Arabidopsis thaliana* plants were grown in a growth chamber on *Jiffy*-7® peat pellets (44x42 mm, 100% peat, http://www.jiffypot.com) at 22°C, under short-day conditions (9h light, 200 µmol/m²/s) with 70-80% relative humidity. Arabidopsis mutant lines *drn-1 drnl-1* (*33*), *cuc2-1* (*41*) (*42*), *cuc2-3* (*43*), *ost2-2D* (*37*), *clf-28* (SALK_139371) and *clf-29* (SALK_021003) mutants (*44, 45*) and transgenic line *CUC2g-m4* overexpressing *CUC2* (*30*) were previously described. The 366 homozygous EMS mutant (HEM) lines were obtained from the Versailles Arabidopsis Stock Center {Capilla-Perez, 2018 #8762, https://publiclines.versailles.inrae.fr/catalogue/hem, data S1}. Cauliflower plants (*Brassica oleracea* cv. *botrytis* var. *Clovis* F1, Vilmorin) were grown under glasshouse conditions. After inoculation, all plants were placed in a growth chamber at 22°C in short-day conditions (8h light, 190 µmol/m²/s) and 70% relative humidity.

#### Bacterial strains and growth conditions

Strains used in this study are listed in Table S1. *Xcc* strains were grown either in MOKA medium (yeast extract 4 g/L, casamino acids 8 g/L, $K_2HPO_4$ 2 g/L, $MgSO_4.7H_2O$ 0.3 g/L) (*46*) and *Pst* strains in KING B medium (Peptone bacto 20 g/L, glycerol 10 g/L, $K_2HPO_4$ 1.5 g/L, $MgSO_4$ $7H_2O$ 1.5 g/L) (*47*) supplemented or not with 1.5% agar (w/vol) and appropriate antibiotics at 28°C: rifampicin (50 µg/mL), kanamicin 146 (50 µg/mL) and chloramphenicol (10 µg/mL).

#### Bacterial inoculation assays

Bacteria were cultivated overnight at 28°C in MOKA (*Xcc* strains) or KING B (*Pst* strains) liquid medium. On the day of inoculation, bacteria were centrifuged at 6000 g for 10 minutes and washed two to three times in 10 mM $MgCl_2$.

Dip inoculation of the second cauliflower leaf was performed using a *Xcc* suspension at $5.10^7$ CFU/mL. Dip or spray inoculations were performed on entire 4-5-week-old Arabidopsis rosettes with bacterial suspension at $5.10^6$ CFU/mL and $2,5.10^6$ CFU/mL for *Xcc* and *Pst*, respectively. Spray was performed by saturating the adaxial leaf surface with a spray bottle. For dip inoculation, the cauliflower leaf or the entire Arabidopsis rosette were dipped into the inoculum for 30 seconds. Plants were left to dry before returning to the growth chamber. Mesophyll infection was achieved by infiltration of a bacterial suspension at $5.10^4$ CFU/mL. All inoculations were carried out in the morning.

In order to manipulate ambient humidity, plants were kept noncovered or covered by a saran wrap or hard transparent cover. Saran wrap was eventually lacerated and referred to as a partially covered condition. Guttation inhibition was achieved by applying silicone grease to the margins of the leaves with a spatula 24 hours prior to inoculation.

In order to determine bacterial populations, spray-inoculated rosettes were harvested, weighed and ground in 10 mM $MgCl_2$. For infiltrated and dip-inoculated plants, one (Arabidopsis) or four (Cauliflower) 6-mm diameter leaf discs were excised per leaf. Serial dilutions were deposited on MOKA (*Xcc*) or KING B (*Pst*) plates supplemented with appropriate antibiotics and 30 µg/mL pimaricin. After incubation at 28°C for 40 hours (*Pst*) or 60 hours (*Xcc*), colonies were counted and bacterial populations expressed as CFU (colony-forming units) per g fresh weight or square cm. Each experiment was repeated at least three times. HEM lines were phenotyped using three independent rosettes. For other Arabidopsis pathoassays, each biological repetition included three leaves from four plants. For cauliflower, each biological repetition included one leaf from four plants.

#### Determination of hydathode numbers and serration

In order to accurately determine hydathode numbers per leaf, leaves were clarified with ethanol and observed under an optical microscope. A single measure was conducted on 2,5-week-old HEM mutant plants to count hydathodes based on three to seven leaves from rank three. For *cuc2* and *drn-1 drnl-1* mutants or *CUC2* transgenics and their wild-types, six-week-old plants were used and the count was performed on 9 to 12 plants in leaf ranks 3 and 5. For serration observations, five-week-old plants were used.

**Induction and observation of water-soaking**

Six-week-old plants were covered overnight under saturating humidity to promote guttation. Three plants per genotype were collected at the end of the night period. For each of the biological triplicates, three leaves were selected and photographed on the abaxial face.

**Visualization of ß-glucuronidase activity *in planta***

Leaf tissues infected with *Xcc* 8004::*GUS-GFP* derivatives were vacuum infiltrated in 50 mM sodium phosphate buffer, pH 7.2, 1 mM 5-bromo-4-chloro-3-indolyl β-D-glucuronide, 0.2% Triton X-100 [v/v], 2 mM potassium ferricyanide [$K_3FeCN_6$], 2 mM potassium ferrocyanide [$K_4FeCN_6$]) and incubated in the dark at room temperature for 24 hours. Samples were cleared in successive baths of 80% ethanol at room temperature until full destaining of the chlorophyll.

**Confocal laser scanning microscopy**

Fluorescence was observed with a confocal laser scanning microscope (TCS SP8; Leica, Germany) using a x63 oil immersion objective lens. UV autofluorescence was excited with a 405nm laser line and detected in the 415/470 nm emission range. GFP fluorescence and chloroplast autofluorescence were excited with the 488 nm laser line and detected in the 500–550/ 680–720 nm emission range, respectively. Images were analyzed using LAS X software (version 3.5, Leica).

**Data processing, representation and statistical analyses**

The data were processed and analyzed using R (version 4.3.2; https://www.r-project.org). All boxplots and scatterplots were generated using the *ggplot2* package. Statistical analyses were performed as indicated in the respective figure legends, including the Kruskal–Wallis test (*agricolae* package) followed by Dunn's post hoc test for pairwise comparisons with Benjamini–Hochberg (BH) correction (*FSA* package), along with Spearman correlation tests (*stats* package). Linear models and linear mixed-effects models to account for random effects (*lme4* package) were also applied, with Tukey- or Sidak-adjusted post hoc comparisons as appropriate. Experiments were independently reproduced at least three times.

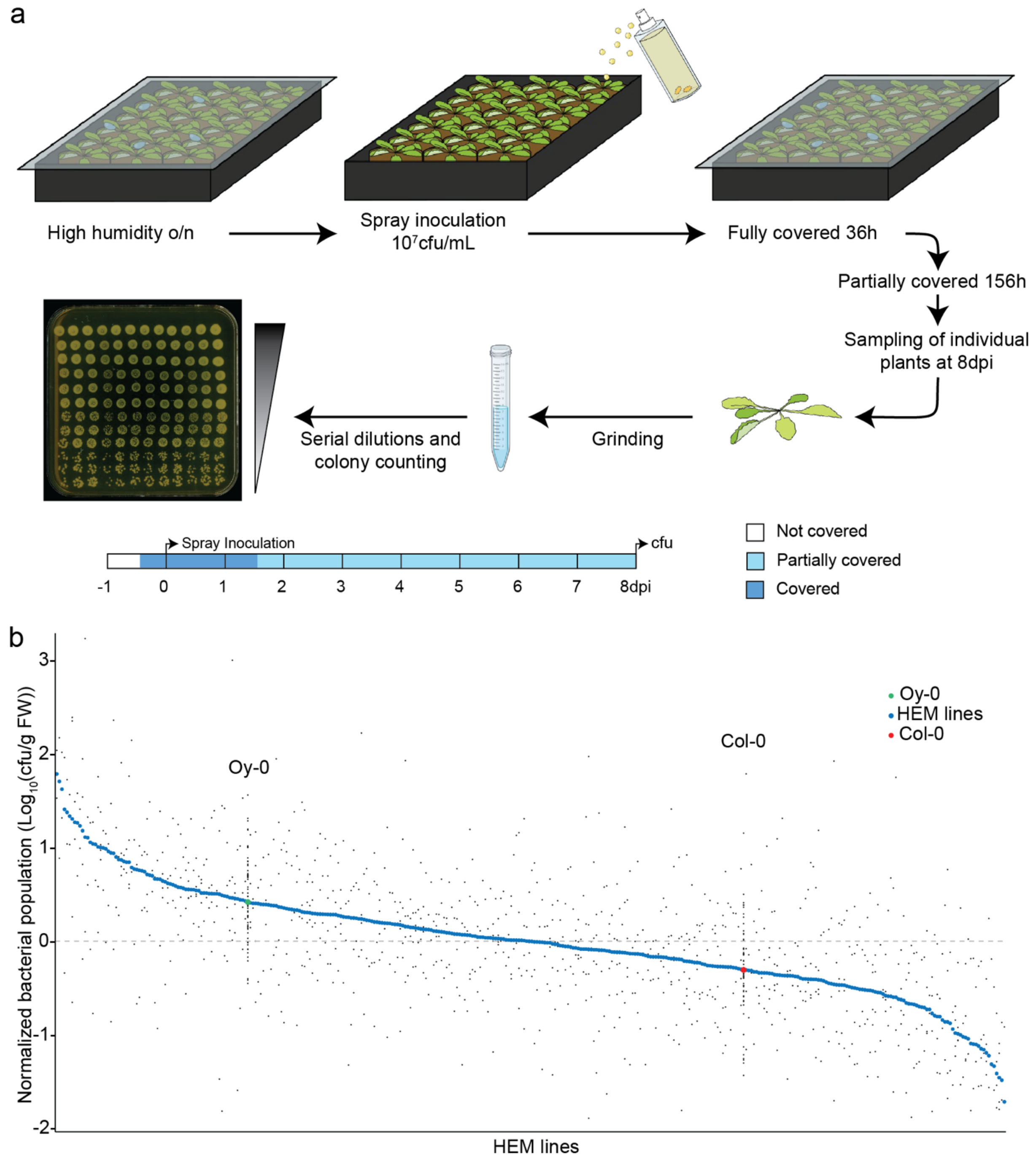


**Fig. S1. Screening of 366 HEM Arabidopsis mutants for their susceptibility to virulent *Xcc* upon spray inoculation and hydathode-mediated infection. (a)** Experimental procedure for efficient and reproducible hydathode-mediated infection of Arabidopsis leaves. At eight days post spray inoculation, total bacterial population is determined on the whole rosette by grinding, cfu counting and normalization to the fresh weight. **(b)** Median *Xcc* 8004Δ*avrAC* populations were determined in a total of 366 HEM lines split in multiple experimental blocks, each including wild-type Col-0 and Oy-0 plants. Each of the three biological repetitions included a single plant per genotype. Individual values were normalized between the different experimental blocks. Colored points represent the median value for each genotype. Black points represent individual values. The Col-0 accession corresponds to the non-mutagenized parental HEM line. The Oy-0 accession is more susceptible to *Xcc* by hydathode infection.

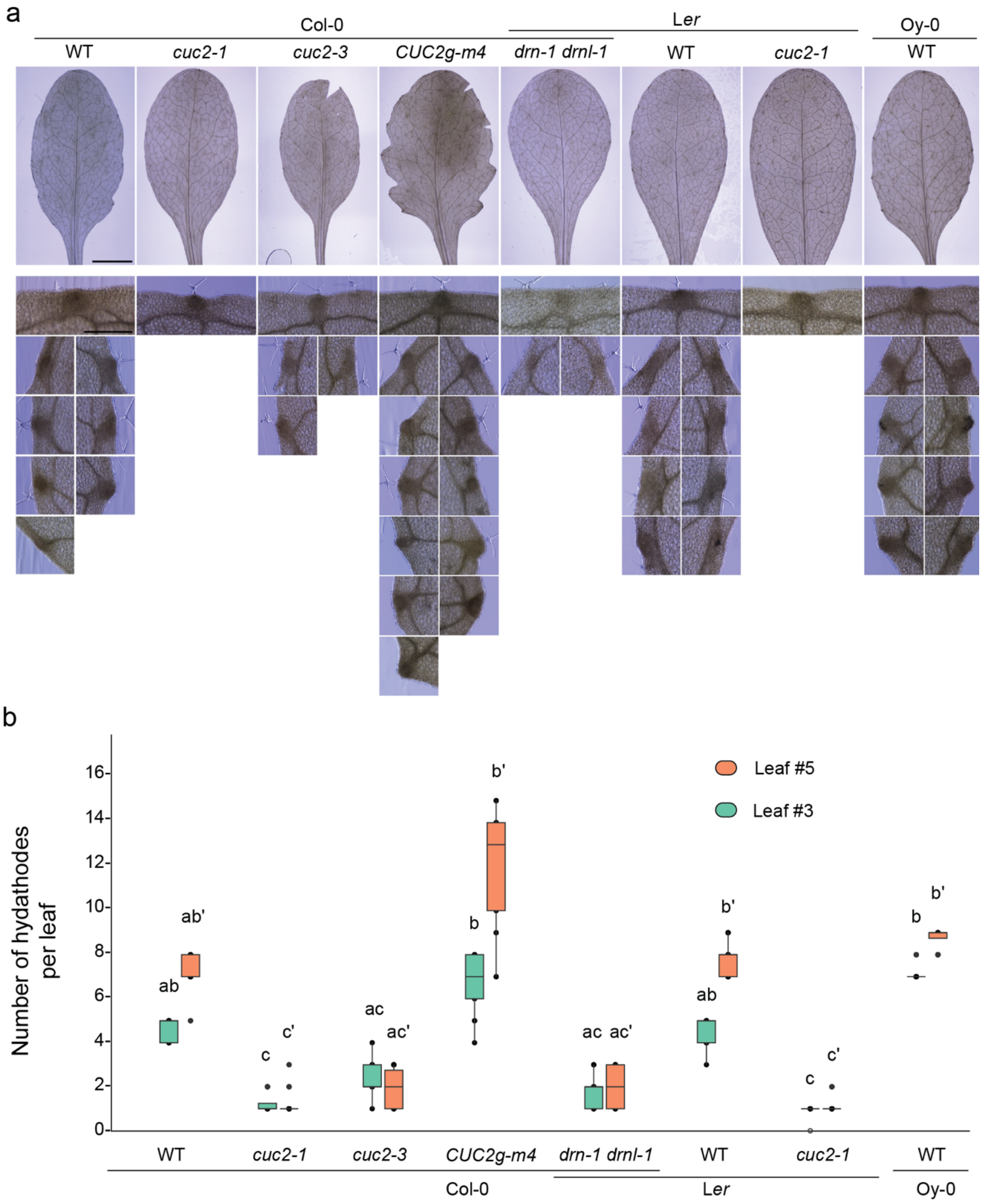


**Fig. S2. Observation and counting of hydathodes in leaves of Arabidopsis mutants and transgenics and their corresponding wild-types. (a)** Observation of serration and hydathode localization in the fifth leaf of wild-type, transgenics and mutant accessions of Arabidopsis. The upper line shows the entire leaves, while lower panels show magnification of the apical and marginal hydathodes. A representative leaf is shown for each genotype Scale bar = 5 mm. **(b)** Number of hydathodes per leaf in *A. thaliana* mutants and transgenics on their third and fifth leaves. Letters indicate statistically distinct groups according to a Kruskal-Wallis nonparametric test ($p<0.05$). The apostrophe (‘) refers to the statistical tests for the fifth leaf.

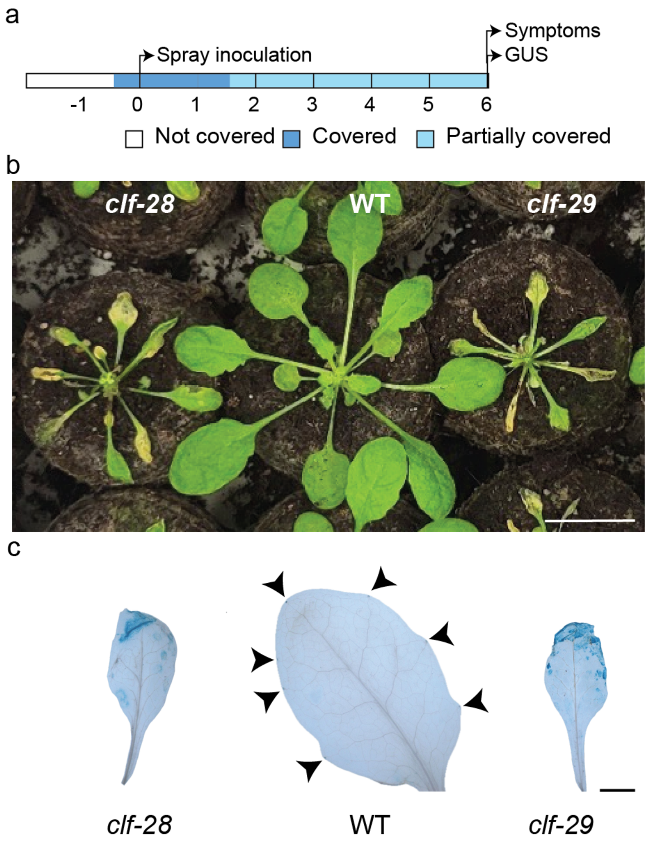


**Fig. S3. *Xcc* can colonize the mesophyll of *clf* mutants. (a)** A timeline depicting the inoculation procedure and sampling is shown. **(b)** Symptoms of wild-type Col-0 plants (WT) and spontaneous water-soaking mutants *clf-28* and *clf-29* six days post spray inoculation of virulent *Xcc* 8004Δ*avrAC::GUS-GFP*. Scale bar = 2 cm. **(c)** Observation of GUS activity produced by *Xcc* 8004*ΔavrAC::GUS-GFP* at six dpi in WT Col-0 plants and *clf-28* and *clf-29* mutants. Arrowheads indicate *Xcc* proliferation in hydathodes. Scale bar = 2 mm.

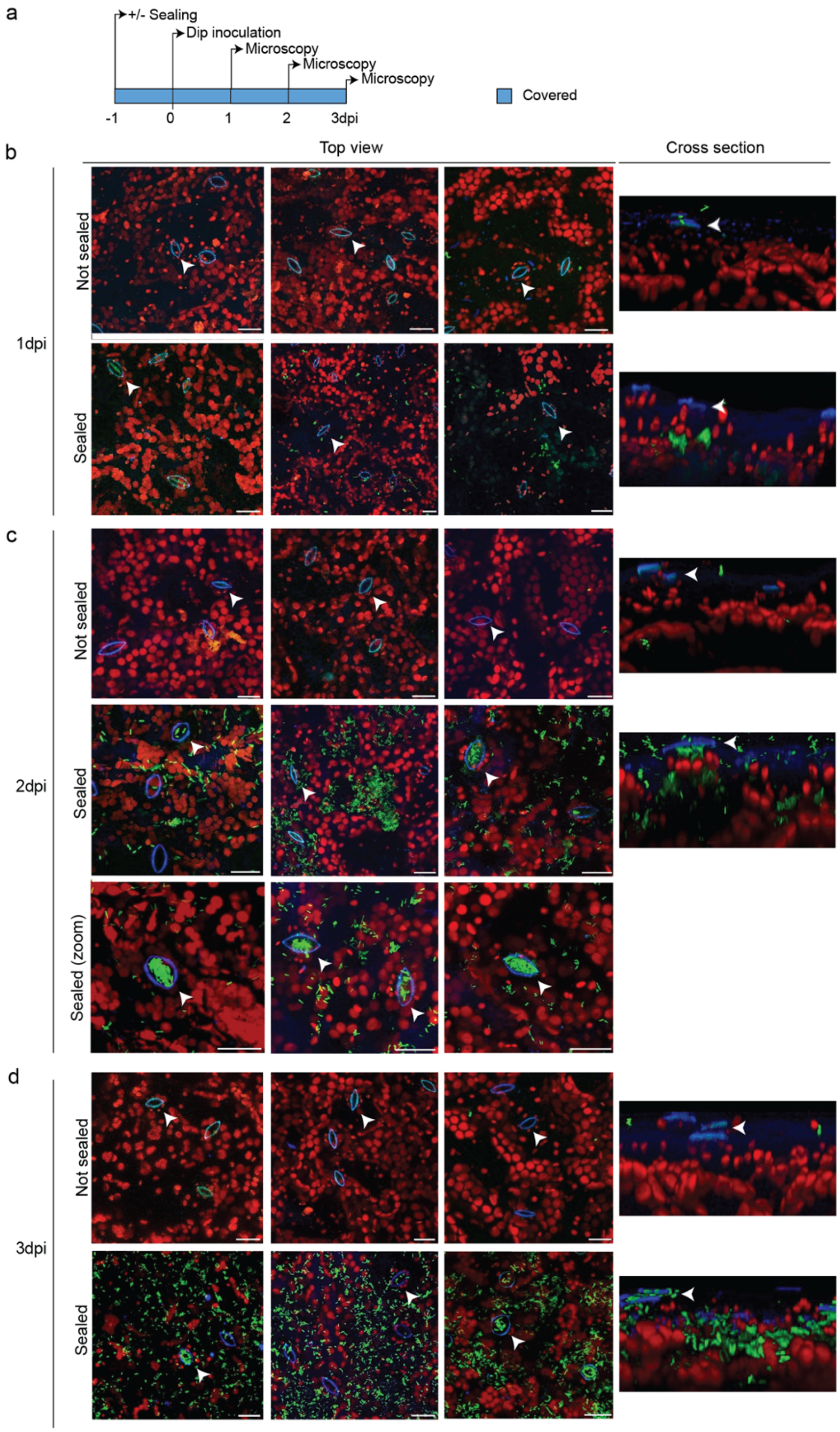


**Fig. S4. Observation of abaxial sides of Col-0 leaves by confocal microscopy at 1, 2 or 3 days post dip inoculation with *Xcc* 8004*ΔavrAC::GUS-GFP* with or without leaf margins sealing.** **(a)** A timeline depicting the inoculation procedure and sampling is shown. **(b-d)** Observation of abaxial sides of Col-0 (WT) leaves by confocal microscopy from above (top view) or cross-section at one (b), two (c) or three (d) days post dip inoculation with *Xcc* 8004*ΔavrAC::GUS-GFP* with or without sealing the margins. Besides autofluorescence (red-chloroplasts and blue-guard cells), GFP fluorescence (green) can be observed in sub-stomatic chambers in sealed conditions only. Arrowheads indicate stomata. Scale bar = 20µm.

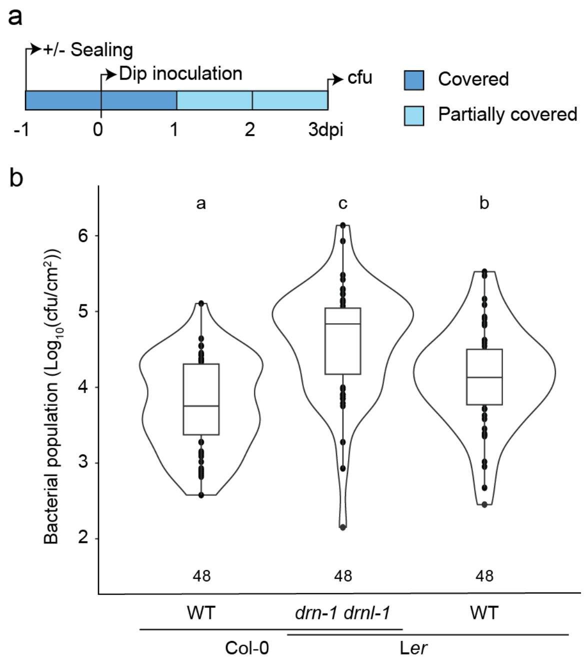


**Fig. S5. Susceptibility of *drn-1 drnl-1* mutant upon dip inoculation with virulent *Xcc 8004ΔavrAC* strain. (a)** A timeline depicting the inoculation procedure and sampling is shown. cfu = colony forming unit, dpi = days post inoculation. **(b)** Boxplot representation of *Xcc* 8004*ΔavrAC* populations in *A. thaliana* mutant and corresponding wild types at three dpi. Letters indicate statistically distinct groups according to a linear model with adjusted comparison ($p<0.05$). The number of leaves sampled is indicated.

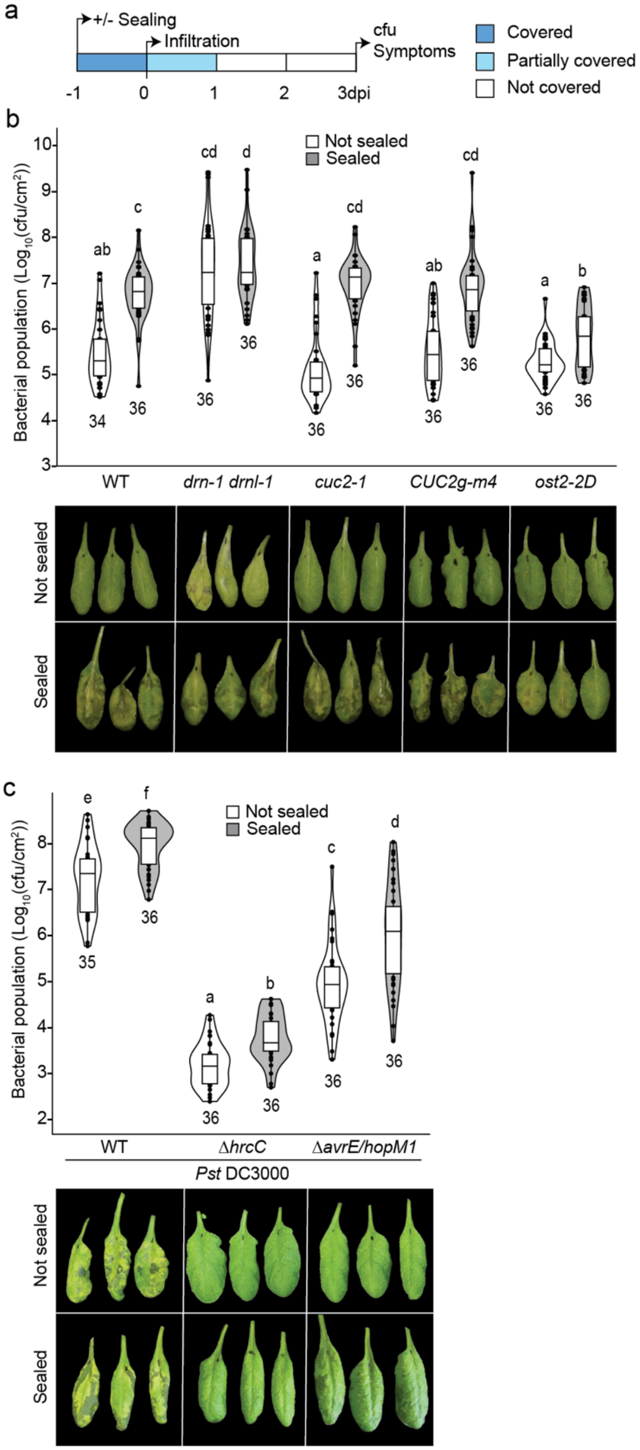


**Fig. S6. Water-soaking in sealed condition promotes bacterial population after infiltration in Arabidopsis Col-0 leaves. (a)** A timeline depicting the inoculation procedure and sampling is shown. **(b)** Boxplot representation of population of virulent *Xcc* 8004Δ*avrAC* in sealed or nonsealed leaves three days post inoculation by infiltration. **(c)** Boxplot representation of population of *Pst* DC3000, *Pst* DC3000*ΔhrcC* or *Pst* DC3000*ΔavrE/hopM1* in sealed or nonsealed leaves 3 days post inoculation by infiltration. Letters indicate statistically distinct groups according to a linear (b) or linear mixed-effects model (c) with adjusted comparison ($p<0.05$). The number of leaves sampled is indicated. Representative images of symptoms are shown.

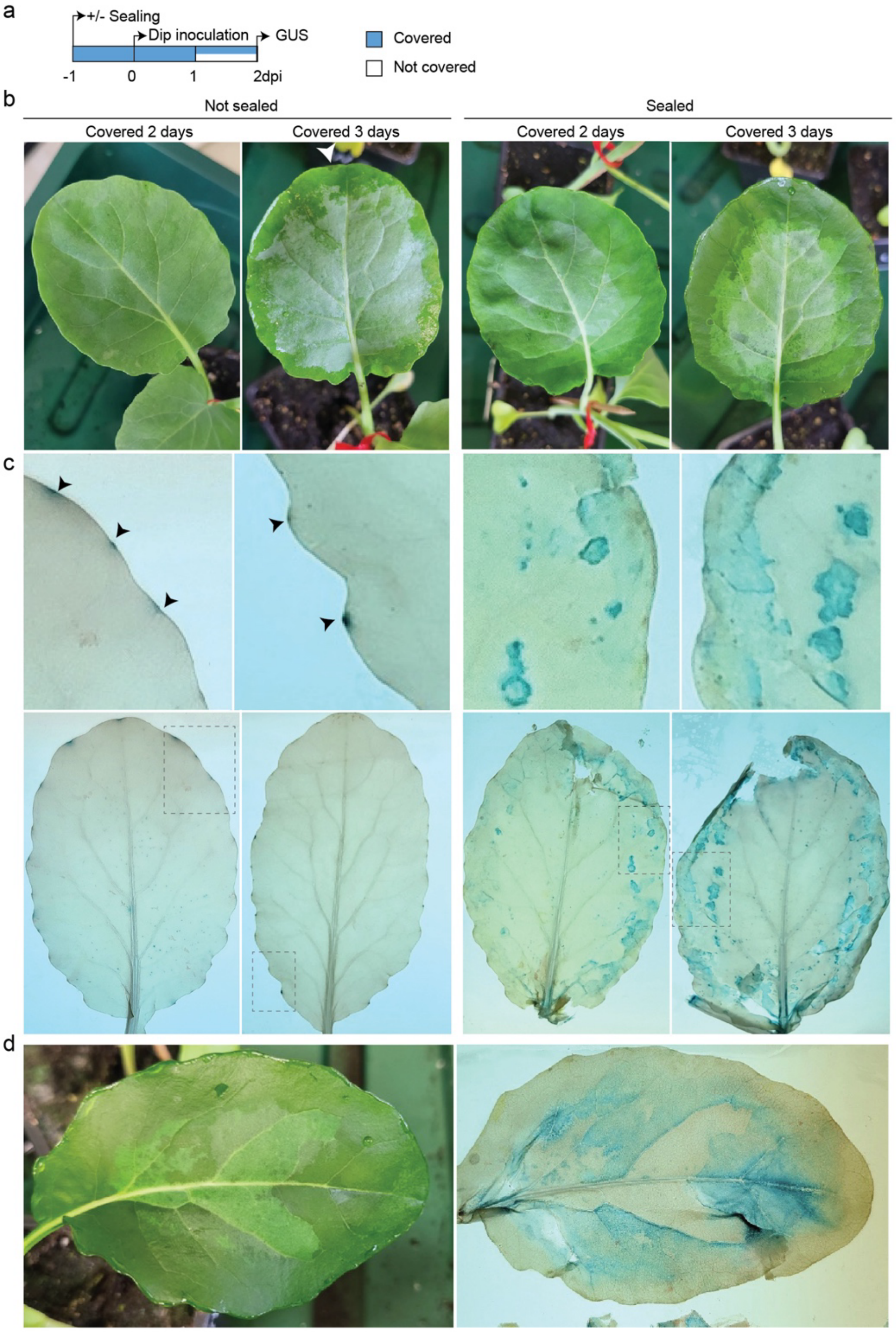


**Fig. S7. Inhibition of guttation induces water-soaking and mesophyll infection in cauliflower.** **(a)** A timeline depicting the inoculation procedure and sampling is shown. **(b)** Water-soaking of mesophyll (dark green area) is promoted upon inhibition of guttation using silicon grease (sealed) and high humidity conditions. Arrowhead indicates the only water-soaked area observed in nonsealed leaves: other wet areas correspond to water present on the leaf surface. **(c)** Observation of GUS activity produced by *Xcc* 8004*::GUS-GFP* strains on cauliflower leaves after two dpi. Bacteria were dip-inoculated one day after sealing leaf margins with silicon grease. Close-ups (top panels) extracted from entire leaves (bottom panels) show the infection of hydathodes in nonsealed leaves (Arrowheads) or of the mesophyll in sealed leaves. **(d)** At two dpi, GUS activity of *Xcc* 8004*::GUS-GFP* strain (right) overlaps with water-soaked areas (left) in a sealed leaf kept under saturating humidity.

**Table S1. List of strains used in this study**

| Strain | Description | Reference or source |
|---|---|---|
| *Xanthomonas campestris* pv. *campestris* | | |
| 8000 | Wild-type strain, harvested on cauliflower, United Kingdom, 1958 | (*38*) |
| 8004 | $Rif^R$-derivative of strain wild-type strain 8000, $Rif^R$ | (*38*) |
| 8004*::GUS-GFP* | 8004 derivative, constitutive expression of GUS and GFP, $Rif^R$ | (*16*) |
| 8004Δ*hrcV* | Avirulent 8004 derivative, carries a *hrcV* deletion, $Rif^R$ | (*16*) |
| 8004Δ*hrcV::GUS-GFP* | Avirulent 8004Δ*hrcV* derivative, constitutive expression of GUS and GFP, $Rif^R$ | (*16*) |
| 8004Δ*avrAC* | 8004 derivative, carries a *avrAC* deletion, $Rif^R$ | (*48*) |
| 8004Δ*avrAC::GUS-GFP* | 8004Δ*avrAC* derivative, constitutive expression of GUS and GFP, $Rif^R$ | (*16*) |
| *Pseudomonas syringae* pv. *tomato* | | |
| DC3000 | Wild-type, $Rif^R$ | (*49*) |
| DC3000Δ*hrcC* | DC3000 derivative, carries a *hrcC* deletion, $Rif^R$ and $Cm^R$ | (*50*) |
| DC3000Δ*avrE1/hopM1* | DC3000 derivative, carries deletions of *avrE* and *hopM1*, $Rif^R$ and $Kan^R$ | (*9*) |

Antibiotic resistances: Kan, kanamycin; Rif, rifampicin; Cm, chloramphenicol.
